# NELFA-mediated promoter-proximal pausing restrains YAP-driven transcription and shapes context-dependent outcomes in breast cancer

**DOI:** 10.64898/2025.12.29.696819

**Authors:** Bhavesh Vasave, Aadya Atreya, Rishabh Kulkarni, Sanket Nagarkar, Khushi Salvi, C B Koppiker, L S Shashidhara, Madhura Kulkarni

**Author notes:** Corresponding Author: Madhura Kulkarni, PhD, Translational Research lead and DBT-Ramalingaswami Fellow, Centre for Translational Cancer Research. Indian Institute of Science Education and Research (IISER) Pune and Prashanti Cancer Care Mission, Pune, B-403, Third Floor, Main Building, Dr Homi Bhabha Road, Pashan, Pune 411 008 Maharashtra INDIA.

## Abstract

Overexpression of YAP is associated with tumor progression in multiple malignancies. YAP is the key transcriptional effector of Hippo signaling: a conserved pathway from *Drosophila* (Yki) to humans (YAP). In a *Drosophila* screen, we identified NELFA, a core component of the promoter-proximal pausing (PPP) complex, as a suppressor of Yki-driven hyperproliferation. We investigated whether PPP–YAP regulatory interaction is conserved in mammalian system. In HEK293T and MDA-MB-231 cells, we demonstrate that NELFA depletion amplifies YAP-driven transcription. Whole-transcriptome analysis of MDA-MB-231 cells revealed widespread reprogramming upon NELFA loss, with strong enrichment of YAP signatures, EMT and TGF-β transcriptional networks—indicating that NELFA regulates a broad YAP-centered gene regulatory module. Clinical analysis of two independent breast cancer cohorts revealed a context-dependent role for NELFA. Low NELFA expression predicted poor disease-free survival in YAP-high tumors—particularly in triple-negative breast cancer (TNBC)—suggesting a tumor-suppressive role. While High NELFA expression correlated with poorer overall survival in all the subtypes. These findings confirm that NELFA’s role in YAP-driven transcription is conserved in mammalian system and it manifests in a context-dependent manner in the breast cancer.

## Introduction

The YES Associated Protein (YAP) is an oncoprotein involved in multiple solid tumor malignancies like breast, lung, liver, etc. [1]. YAP binds to the TEAD family of proteins as a transcriptional co-activator for its transcriptional activity [2,3]. And YAP-TEAD interaction is essential for the tumorigenic gene expression [2] involved in proliferation, EMT, migration and invasion [4]. The Drosophila ortholog of YAP, known as Yorkie (Yki), also regulates genes involved in cell growth and survival, including Diap1, dMyc, and bantam [5]. A genetic screen conducted to identify tumor suppressors that regulate Yki-driven hyperproliferation; a surrogate for tumor proliferation, revealed NELFA, a component of the promoter proximal pausing complex, as one of the significant tumor suppressors [6].

Promoter-proximal pausing (PPP) is a key regulatory checkpoint in transcription where RNA Polymerase II (Pol II) transiently halts shortly after initiation [7]. This pause is stabilized by the NELF complex (NELFA, B, C/D, E), which binds the Pol II–Spt5 interface and prevents premature elongation by distorting the active site and blocking TFIIS-mediated rescue [8,9]. Involvement of this PPP complex in mediating Yki-driven hyperproliferation was confirmed in *Drosophilla*, where knocking down individual components of PPP complex enhanced Yki-driven tumorigenesis [5].

Because both YAP-Hippo signalling and the PPP machinery are evolutionarily conserved, these findings suggested a potential regulatory interface between NELF-mediated pausing and YAP-driven transcription in mammalian systems. Breast cancer provides an appropriate context in which to examine this interaction, as the oncogenic role of YAP in mediating proliferation, epithelial-to-mesenchymal transition, therapeutic resistance, and metastatic dissemination is well established in this context [2,10]

In this study, we investigated whether NELFA modulates YAP-dependent transcription in mammalian breast cancer cells and evaluated the broader biological and clinical implications of NELFA and YAP axis. We assessed the effects of NELFA depletion on YAP target-gene expression using transcriptomic profiling and examined the impact of the NELFA–YAP association in two patient cohorts: the TCGA BRCA dataset and an independent breast cancer cohort from our biobank. Together, these complementary approaches reveal a context-dependent role for NELFA in shaping YAP-driven transcriptional programs and influencing breast cancer patient outcomes.

## Methods and methodology

### Cell culture

HEK293T; MDA-MD-231 and SKBR3 cell lines were used in the study. HEK293T and MDA_MB-231 cells were a gift from Prof. Ito at CSI, Singapore whereas SKBR3 cells were obtained from NCCS (National Centre for Cell Science). HEK293T and MDA-MB-231 cell lines were cultured in (DMEM) Dulbecco’s Modified Eagle Medium High glucose (HIMEDIA-#AL066A) supplemented with 1x sodium pyruvate (GIBCO-Cat. No. 1136007), 10% FBS (Fetal Bovine Serum, qualified, Brazil, GIBCO – Cat. No. 10270106) and 1% Penicillin Streptomycin (Penicillin-Streptomycin, Sigma-Aldrich, Cat. No. P4333) in standard conditions incubated at 37° C and 5% CO2. SKBR3 cell lines were cultured in McCoy’s 5A supplemented with 10% FBS and 1% Penicillin-Streptomycin.

### si-NELFA and YAP overexpression in HEK293T

HEK293T cells were stably transfected with pMSCV empty vector or pMSCV with Flag-YAP S127A/S397A; a gift from Prof Stephen Cohen, as reported in Nguyen et al., 2014. SMARTpool siRNAs for scramble, NELFA, HEXIM1, HEXIM2, and MEPCE were obtained from Sigma-Aldrich, and were individually transfected into HEK293T cells with stable expression of YAP-S127A/S397A. Knockdown was confirmed in 48 to 72 hrs by RT-PCR.

### si-NELFA and si-YAP in MDA-MB-231

siRNA against NELF-A (ON-TARGETplus Human NELFA siRNA smartpool #L 012156-00-0005) YAP (ON-TARGETplus Human YAP1 siRNAsmartpool #L-012200 00-0005) and control siRNA (Non-targeting Pool #D-001810-10-05) were ordered from Dharmacon. MDA-MB-231 cell line was transfected with 50-100nM siRNA concentration using DharmaFECT Transfection Reagent (T-2001-02) according to the manufacturer’s protocol. Cells were harvested at 80% confluency, and RNA was extracted at two time points: 48 hours and 72 hours after transfection.

### RNA extraction and RT-qPCR

RNA was isolated using TRIzol (Invitrogen, Cat. No. 15596026) extraction protocol. Reverse transcription was performed using iScript cDNA Synthesis Kit (BioRad, #1708891) according to the manufacturer’s protocol. RT-PCR was then performed using the pre-amplified cDNA using the iTaq Universal SYBR Green Supermix (BioRad, #1725121) kit according to the manufacturer’s protocol. Normalization of RT-PCR was computed using Ct values with respect to the housekeeping gene GAPDH. The list of primers used is provided in Table 1.

### Immunoblotting for knock-down confirmation

Whole cell lysates from the transfected cell lines were extracted from the confluent cell cultures using a modified RIPA buffer prepared in-house (20 mM Tris-HCl, pH 8.0, 420 mM NaCl, 10% Glycerol, 0.5% NP-40, 0.1 mM EDTA, with 1 mM DTT, 10 mM PMSF, and 20 mM protease inhibitor). The protein concentration was estimated using the Bradford assay. Equal concentrations (2ug/ul) of proteins were fractionated by SDS-PAGE and transferred onto PVDF membrane. Protein blots were probed overnight with primary antibodies diluted in 10% milk at 4°C. Subsequently, the blots were incubated with HRP-tagged secondary antibodies and then imaged using the ImageQuant LAS 4000 biomolecular imager, where bands were detected via chemiluminescence. The intensity of bands was quantified using ImageJ software. The list of antibodies used is provided in Table 2.

### Whole Transcriptome RNAseq

Total RNA was isolated from MDA-MB-231 cells at 72 hours post–siRNA transfection, obtained from three independent biological replicates. RNA samples were quantified and adjusted to 50 ng/µL. RNA integrity and purity were assessed using the Qubit RNA BR Assay (Invitrogen, Cat# Q10211) and the Agilent TapeStation with RNA ScreenTapes (Agilent, Cat# 5067-5576), and only samples with RIN ≥ 7 were used. Strand-specific total RNA libraries were prepared using the KAPA RNA HyperPrep Kit with rRNA depletion and sequenced on the Illumina NovaSeq X Plus platform to generate 150 bp paired-end reads.

The resulting raw reads were then processed through a standard RNA-seq analysis pipeline. High-quality reads were aligned to the human reference genome (GRCh38, Ensembl release 87) using STAR with the two-pass mapping strategy [11]. Quality control of the aligned data was further assessed using RNA-SeQC [12], RSeQC [13], and MultiQC [14]. Gene-level quantification was performed using featureCounts [15]. Transcript abundances were estimated in FPKM and TPM. Differential gene expression analysis was performed using the DESeq2 package [16].

### Gene-set Enrichment Analysis

Gene set enrichment analysis (GSEA) was performed in R using the **clusterProfiler** [17], **fgsea** [18], and **msigdbr** package [19]. All protein-coding genes (based on Ensembl biotype annotation) that were significantly differentially expressed (nominal *p* < 0.05) in siNELFA vs siControl, siYAP vs siControl, and siNELFA + siYAP vs siControl were taken as input. Ensemble gene IDs were converted to HGNC gene symbols using the org.Hs.eg.db package [20]. Genes were ranked by log2 fold change, and enrichment was tested against the MSigDB v7.5.1 collections: C6 Oncogenic Signatures and H Hallmark Gene Sets [21]. Visualization of enriched pathways was performed using dot plots, which displayed Normalized Enrichment Scores (NES), −log₁₀ (nominal *p*-values), and gene set sizes. Venn diagram was constructed to visualize overlapping gene sets using the ggVennDiagram package with labelled set sizes and intersection counts [22]. Distribution of genes across the four mutually exclusive regulatory categories was visualized via an UpSet plot generated using the ComplexUpset package in R [23].

### Transcription Factor (TF) Enrichment Analysis

Transcription factor binding site enrichment analysis was performed using the **Enrichr** platform [24] via its web interface (https://maayanlab.cloud/Enrichr/). The input comprised all significantly differentially expressed protein-coding genes (p < 0.05) from the siNELFA vs siControl comparison, without applying a log₂ fold change cutoff. Four TF-related gene set libraries were queried for enrichment: ChEA 2022, ENCODE TF ChIP-seq 2015, ENCODE and ChEA Consensus TFs from ChIP-X, and TF Perturbations Followed by Expression. For each enriched term, the overlap count, nominal p-value, and Enrichr’s Combined Score were recorded. Visualization was performed in R using **ggplot2 [25]** and with dot plots displaying enrichment terms (Y-axis), number of overlapping target genes between the siNELFA vs siControl and the corresponding TF target gene list (X-axis), –log₁₀(p) as bubble color, and Odds ratio (effect size) as dot size. Significant terms appearing across multiple human libraries were selected and ranked according to the absolute number of overlapping gene targets.

### Data and Code Availability

All RNA-sequencing data generated in this study have been deposited in the NCBI Gene Expression Omnibus (GEO) under accession GSE311396 (https://www.ncbi.nlm.nih.gov/geo/query/acc.cgi?acc=GSE311396). The corresponding raw FASTQ files are available in the NCBI Sequence Read Archive (SRA) under BioProject PRJNA1366098, with associated BioSample accessions SAMN53303610–SAMN53303619 (https://www.ncbi.nlm.nih.gov/bioproject/PRJNA1366098).

All scripts used for data processing, statistical analysis, and figure generation are publicly available at GitHub (https://github.com/tmemklab/siRNA-NELF-A-MDA-MB-231).

### Patient sample procurement and ethics

Primary breast tumor samples (formalin-fixed paraffin-embedded, FFPE), along with their associated de-identified patient metadata, were received from the Prashanti Cancer Care Mission (PCCM) Biobank, with appropriate patient consent and ethical approval (dated 1^st^ November 2022). Seventy-five patients who were diagnosed and underwent treatment from 2010 up to 2020 were included in the study cohort.

Molecular subtypes of these breast tumors were evaluated by determining ER/PR expression and HER2 scores using immunohistochemical analysis and FISH reports from a recognized pathology laboratory. Samples with more than 1% ER expression were taken as ER+. Samples with 0, 1+ or 2+ IHC scores and a negative FISH report were taken as negative for HER2 and positive or negative for PR, while samples with 2+ or 3+ IHC scores but positive for FISH were categorized as HER2+ and negative for less than 1% ER expression irrespective of PR expression. Samples with less than 1% Er and PR expression each, and IHC scores of 0, 1+ or 2+ with a negative FISH report for HER2 were categorized as triple negative.

Following the guidelines provided by the National Comprehensive Cancer Network (NCCN), NACT and ACT treatment were administered to the patients. Twenty-nine patients underwent NACT, for whom response to treatment was determined by comparing cT and cN against ypTypN. yPT0/Tis0, ypN0 status as considered as a pathological Complete Response (pCR) and the rest were categorized as residual disease (RD).

### Immunohistochemistry for YAP

75 FFPE primary tumor samples were sectioned into 3 µm sections using Leica Microtome RM2255 on positively charged hydrophobic slides (PathnSitu, #PS011-72). Tissue slides were deparaffinized, cleaned, and processed for immunohistochemistry using UltraVision Quanto Detection System HRP DAB (Epredia, TL-125-QHD) according to the manufacturer’s protocol. Antigen retrieval was performed using a TRIS-EDTA buffer at pH 6.0 for YAP, which had already been standardized in the lab. Primary antibody treatment for YAP (AbCam, #ab52771) was performed at a 1:200 dilution, and the slides were incubated overnight at 4°C. Tissue samples were stained and scored for YAP expression as reported (Bhardia et al, in revision).

### Immunohistochemistry for NELFA and NELFB

To optimize immunohistochemistry (IHC) conditions for NELFA (Santa Cruz, #sc-365004) and NELFB (Abcam, ab167401) antibodies, we tested epitope retrieval at pH levels of 6, 8, and 9 on high-tumor-content tissues using a 1:50 dilution of the antibody. While pH 6 and 8 yielded variable staining, pH 9 buffer consistently produced the most uniform nuclear staining. Further testing with multiple dilutions of the antibodies revealed that a 1:100 dilution was the optimal setting for IHC (Supplementary Figure 5). In the preliminary assessment, we did not observe any differences in staining patterns for NELFA and NELFB. Since the NELFA subunit of the PPP complex interacts directly with the transcriptional machinery [26,27] we proceeded with NELFA IHC for further studies.

### Imaging and scoring of stained slides

All the immunohistochemistry slides were imaged at 400X by OptraScan using OS-15 bright field digital scanner. Images were checked for focus and quality, converted to TIFF format, and scale bars were added using Image Viewer Version 2.0.4 software provided by OptraScan. Slides were scored for YAP and NELFA percent expression and the intensity by a certified pathologist. Percent scores were binned and multiplied by intensity scores to generate a composite score. The composite scores were used for the ROC curve against DFS in months using the IBS SPSS software.

### Clinicopathological and Survival Analysis

The distribution of clinicopathological characteristics within the cohort and breast cancer subtypes was analyzed using a 2 × 3 (or 4 × 3, in the case of tumor size) Chi-square contingency test, and the results were computed using GraphPad Prism v.8.

Survival outcome analysis was carried out using the Kaplan-Meier method. Overall survival (OS) and disease-free survival (DFS) analyses were performed for a five-year follow-up. Overall survival (OS) was defined as the time in months from diagnosis to death or the last follow-up date, and Disease-free survival (DFS) was defined as the time in months from the date of surgery to the date of recurrence or the last follow-up date. Survival probabilities were calculated using the Log-rank test statistics in GraphPad Prism.

### BRCA cohort analysis from the TCGA dataset

RNA sequencing and corresponding clinical data were extracted using R (version 4.0.0). The github link has the script used-https://github.com/tmemklab/tcga_data_download/blob/main/run_tcga_biolinks.Rmd. The dataset included patient sample information, associated clinical metadata, and expression values in FPKM, TPM, and raw count formats, along with HGNC gene identifiers. All data files were downloaded in CSV format for downstream analysis.

Relevant information was curated using Microsoft Excel to isolate the desired subsets. Only invasive ductal carcinoma (IDC) samples, as described in [28] were selected for further analysis. For each patient sample, TPM values, overall survival (OS), and disease-free survival (DFS) data (in months) were retrieved.

To categorize gene expression levels, the median TPM value was used as the cutoff to define high and low expression groups. Kaplan–Meier survival analyses were performed using GraphPad Prism (version 8) to generate survival curves based on OS and DFS.

## Results

Promoter proximal pausing (PPP) complex comprises NELF proteins, 7SKsnRNP RNA-Protein subcomplex, and PTEF-b [7,29]. Our previous work in *Drosophila* showed that RNAi-mediated knockdown of 7SKsnRNP components: Bin3 (MePCE ortholog), Hexim (HEXIM1/2 ortholog), and NELF components of the PPP complex enhanced Yki-driven neoplastic transformation of wing imaginal discs [5]. To investigate if the role of the PPP complex in regulating YAP-driven transcription is conserved in the mammalian system as well, we performed siRNA-mediated knockdown of PPP complex components in mammalian cell lines and quantified the impact on YAP-target gene expression.

### NELFA regulates YAP-driven transcription in mammalian cell lines

YAP was stably overexpressed in HEK293T cells, followed by siRNA-mediated depletion of key components of the 7SK snRNP complex (MePCE, HEXIM1/2) or the NELF complex (NELF-A) (Figure S1A–C). RT-qPCR analysis of the direct YAP targets *Cyr61*, *CTGF*, and *ANKRD1* confirmed strong induction upon YAP overexpression. In the YAP-overexpression background, knockdown of MePCE or HEXIM1/2 caused a modest, non-significant reduction in *Cyr61* and *CTGF*, with no effect on *ANKRD1* (Figure 1A–B). In contrast, NELF-A knockdown selectively increased *CTGF* expression, but not *Cyr61* or *ANKRD1* (Figure 1C).

**Figure 1.**
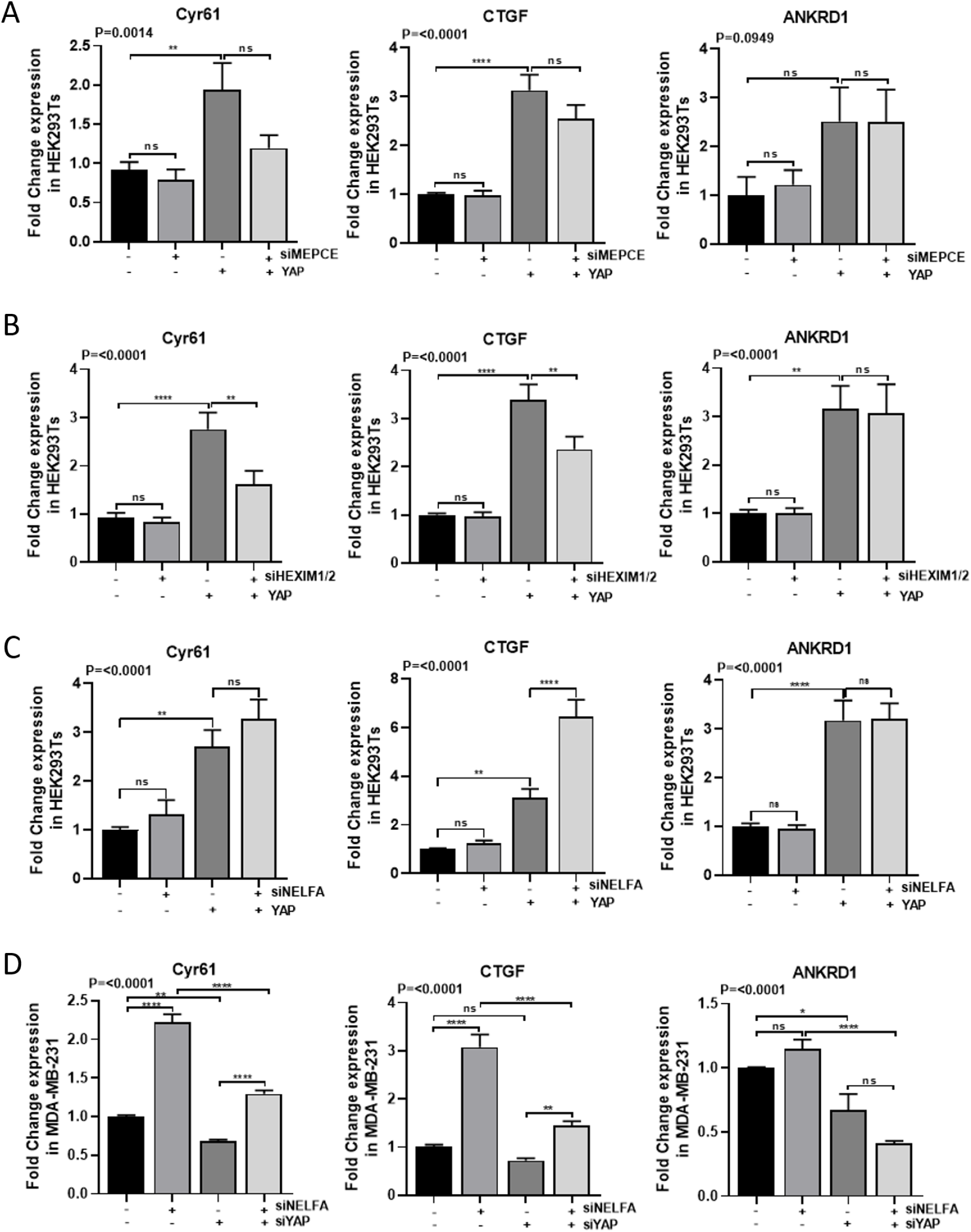
Silencing PPP Components and YAP Alter YAP Target Gene Expression in HEK293Ts and MDA-MB-231 cells. **(A)** Relative mRNA levels of YAP target genes (Cyr61, CTGF, ANKRD1) following siRNA-mediated knockdown of MePCE (7SKsnRNP subunit) in HEK293Ts (N=4). **(B)** Expression of Cyr61, CTGF, and ANKRD1 after HEXIM1/2 knockdown in HEK293Ts (N=4). **(C)** Expression of Cyr61, CTGF, and ANKRD1 following NELFA (NELF complex subunit) knockdown in HEK293Ts (N=4). **(D)** Expression of YAP target genes after NELFA knockdown in MDA-MB-231 cells (N=3). mRNA levels were quantified by RT-qPCR and normalised to GAPDH. Statistical analysis was performed using one-way ANOVA with multiple comparisons; p-values are indicated, with P<0.05 considered significant compared to control.

We further validated the impact of NELF-A knockdown on YAP-target gene expression in MDA-MB-231 breast cancer cells (Figure S1G), which show high endogenous YAP and YAP-target expression (Figure 1D). Strikingly, NELF-A depletion increased *Cyr61* and *CTGF*, but not *ANKRD1*, and this significant upregulation persisted even when YAP was concomitantly knocked down (Figure 1D).

### NELFA CRISPR knockout cells do not survive

To assess the contribution of NELFA and the PPP complex to global and YAP-mediated transcription, we generated CRISPR-based NELFA knockouts in YAP-overexpressing MCF10A and SKBR3 cells. Two NELFA-targeting gRNAs were cloned into the TLCV2 vector, and lentivirus produced in HEK293T cells was used for transduction (Figure S2A). After puromycin selection, doxycycline-induced CRISPR activation was confirmed by GFP expression, followed by FACS isolation of GFP-positive cells (Figure S2B). However, despite two independent attempts, GFP-positive NELFA-knockout cells failed to survive beyond one to two passages, suggesting that NELFA may be essential for cell viability.

### Whole Transcriptome Analysis for NELFA-regulated gene expression

Since CRISPR knock-out generated cells did not survive, global gene expression changes after NELFA knockdown were assessed in MDA-MB-231. Differential gene expression in cell lnes treated with siNELFA vs siControl was analyzed, and Gene-set enrichment analysis (GSEA) was performed [30]. Interestingly, one of the top upregulated gene signatures (NES: 1.68) with high significance (p = 0.003) that showed up is the CORDENONSI_YAP_CONSERVED_SIGNATURE [31] (Figure 2A). Amongst the significantly downregulated pathways, BRCA1-Associated Signature showed the highest enrichment (Figure 2A).

**Figure 2.**
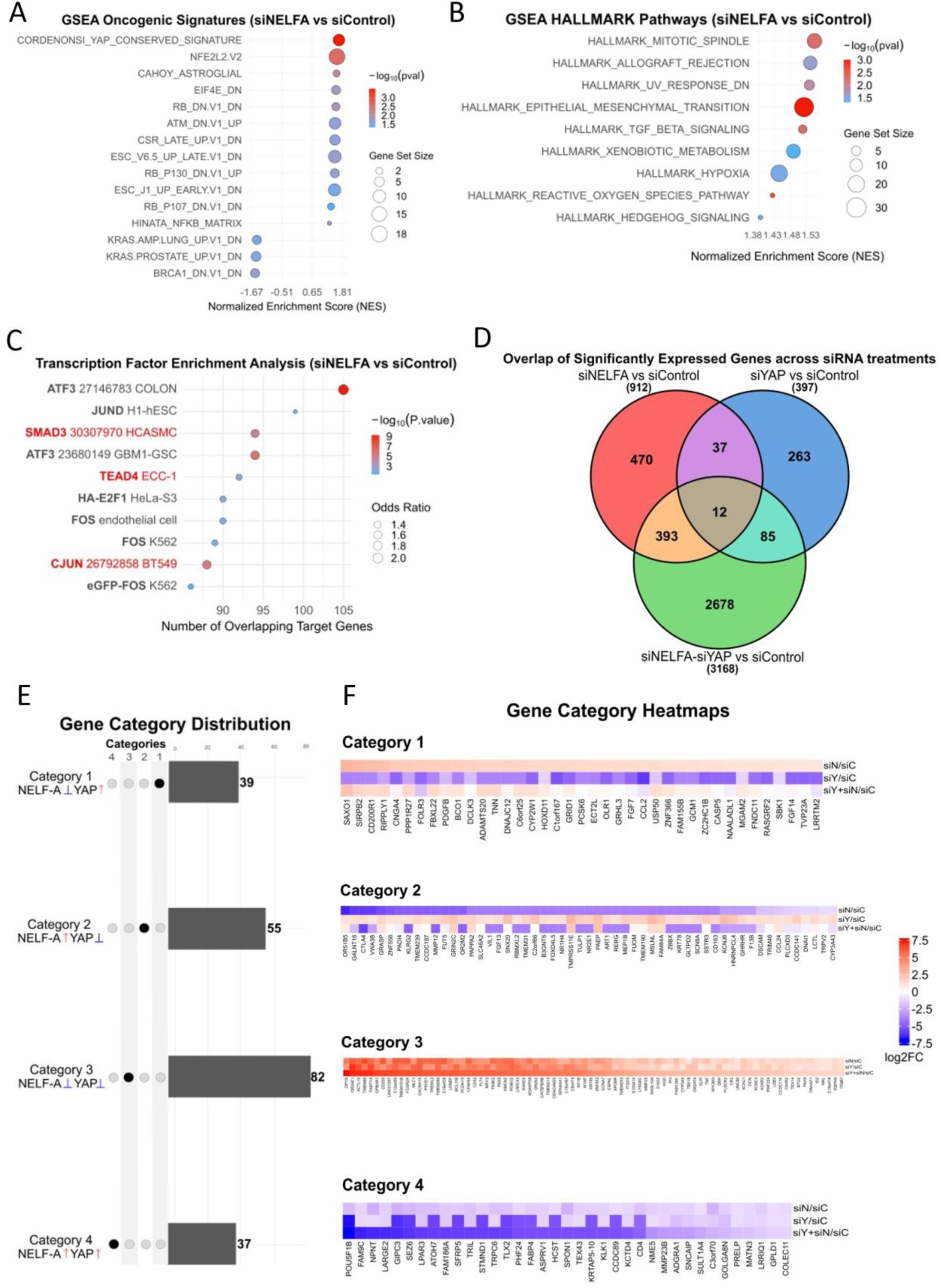
Whole Transcriptome Analysis for NELFA-regulated gene expression. **(A)** Gene Set Enrichment Analysis (GSEA) for MSigDB Oncogenic Signatures (C6) collection of significantly altered genes (p < 0.05) after NELFA knockdown compared to control knockdown. The Normalized Enrichment Score (NES) is plotted on the x-axis, with dot size corresponding to the number of genes overlapping with the ranked list, and color indicating statistical significance (−log₁₀ *p*-value), with red denoting higher significance. **(B)**GSEA Enrichment analysis for MSigDB HALLMARK Pathways. **(C)** Transcription Factor (TF) enrichment results from user-defined gene sets using a combination of perturbation and binding datasets. Y-axis lists the enrichment terms from Enrichr. Each term represents either a TF perturbation experiment (e.g., RNAi, overexpression) from public datasets (like GEO) or a TF binding profile from ENCODE ChIP-seq data (2015). Dot Size reflects the significance of enrichment with the z-score. Dot colour encodes the −log₁₀(p-value) from Fisher’s exact test, with red indicating higher significance. **(D)** Venn diagram showing the overlap of significantly differentially expressed genes (p < 0.05) across siNELFA, siYAP, and siNELFA-siYAP knockdowns compared to siControl. **(E)** Upset Plot showing the distribution of genes across four regulatory categories. Genes were classified based on log2FC≥ 1 thresholds from siNELFA, siYAP, and double knockdown comparisons. Each bar represents the number of genes exclusively assigned to one of the four categories. **(F)** Heatmap of genes grouped into four categories based on log2 fold change

Further, GSEA was also performed on the ranked list of genes using the Hallmark pathways collection to identify broad cancer themes. The most enriched pathway identified was HALLMARK_MITOTOTIC_SPINDLE (NES = 1.56; p= 0.0051) (Figure 2B). Other significantly enriched gene-signatures that showed up were HALLMARK_EPITHELIAL_TO_MESENCHYMAL_TRANSITION (EMT) (NES=1.52 and p=0.0011) and HALLMARK_TGF_BETA_SIGNALING signaling (NES = 1.52, p=0.0046) [21]. Taken together, the results from the Hallmark analysis suggest a model in which NELFA suppresses EMT and its associated invasive and proliferative programs, aligning with the oncogenic functions of YAP.

### Transcription factor target enrichment in NELFA-regulated gene sets

Further, transcription factor (TF) enrichment analysis was performed on all protein-coding genes that were significantly enriched in siNELFA compared to siControl (p < 0.05), to assess the TFs whose transcription may be regulated by NELFA. The top enriched transcription factors ranked by number of overlapping targets included ATF3, JUND, CJUN, SMAD3, TEAD4, and FOS (Figure 2C). Together, the enrichment of TEAD4, SMAD3, and CJUN target genes supports a model in which NELF-A represses a multi-faceted YAP-driven transcriptional network.[1,32]

### Comparison of PPP-regulated genes across studies

Two other reports investigate the effect of PPP on global gene regulation in breast cancer cell lines [33,34]. *Zhang et al.* knock down NELFE in BT549 and MCF7Ras, CRISPR-based knock out of NELFE in SUM159 and MCF7 cell lines , and *Sun et al.* knock down NELFA in the T47D cell line. The DEGs derived from the siNELFA vs. siControl comparison in MDA-MB-231 from this study were compared for overlap with the DEGs regulated by the NELF complex in the other two studies. Pairwise comparison of DEGs across the six breast cancer cell lines revealed little overlap following NELFA or NELFE perturbation in various breast cancer cell lines (Figure S3A).

### NELFA and YAP co-regulated gene sets

To investigate the extent to which NELFA and, thereby, PPP regulate YAP-driven transcription, YAP knockdown was also performed with or without siNELFA in MDA-MB-231. List of DEGs with significant alterations over siControl (p < 0.05) was considered for further analysis. siNELF-A and siYAP knockdown resulted in significant perturbation of 912 and 397 genes, respectively, while the double knockdown perturbed 3168 genes (Figure 2D). However, between the siNELFA and siYAP conditions, only 49 genes were commonly perturbed, and between all three conditions, the overlap showed only 12 common genes (Figure 2D). CTGF, Cyr61, and ANKRD1 exhibited similar expression patterns to those observed by RT-PCR (Figure S3B-D) [36].

The significantly altered genes between siNELFA, siYAP, and siNELFA+siYAP and siControl were compared, revealing four distinct categories of genes co-regulated by YAP and NELFA (Figure 2E and 2F). STRINGdb network enrichment analysis of this group of genes yielded a significant Protein-Protein interaction (PPI) cluster for each category (Figure S3E-G) [35]. The first category comprised 39 genes activated by YAP and suppressed by NELF-A that were upregulated upon depletion of NELF-A and downregulated upon depletion of YAP. Five genes: *CCL2, CD200R1, FGF7, OLR1,* and *PDGFB* from this list showed significant PPI enrichment with p-value = 7.06e-05 (Figure S3E). The second category comprised genes activated by NELFA and suppressed by YAP, which were upregulated upon YAP depletion and downregulated upon NELFA depletion. Significant PPI enrichment for C*TLA4, CD163, MMP12, CCL24,* and *PADI4* was observed (Figure S3F). The third category comprised genes co-repressed by both NELFA and YAP, which were upregulated upon depletion of either factor or further elevated in the double knockdown condition. Network enrichment analysis revealed a highly significant functional network (19 nodes, 29 edges; PPI enrichment p = 1.3e-12), with *TNF* emerging as the central hub (Figure S3G). Finally, the fourth category consisted of genes co-activated by both NELFA and YAP, which were downregulated upon depletion of either factor or most strongly suppressed in the double knockdown condition. In contrast to the other categories, no significant functional clustering of genes was detected in this group.

### Breast cancer cohort characteristics

Our previous *Drosophila* study [5,6] and mammalian cell lines, including HEK293T and MDA-MB-231, demonstrated clear co-regulation of YAP-target genes by NELFA. To investigate whether the YAP-NELFA axis has any implications in breast cancer progression, a cohort of 75 primary breast cancer tumors was assessed for the association between YAP1 and NELFA expression and patient outcomes (Figure S4). The demographic and clinicopathological characteristics of the cohort, according to the molecular subtypes, are presented in Table No. 5. Within the IDC cohort, TNBC reflected a significantly higher proportion of high-grade tumors (73.91% grade III) compared to ER+ and HER2+ (22.6% and 55.6%, respectively). The cohort had a median follow-up of 30 months. Out of 75 patients, 11 patients recurred, and 4 patients died during the five-year follow-up.

**Table 5:** Demographic table of the breast cancer cohort. A cohort of IDC patients grouped according to the molecular subtype, ER+, HER2+ and TNBC subtypes. The distribution of the clinical parameters such as age at diagnosis, menopausal status, tumour grade, radiological and pathological tumour size, stage, LVI, etc are listed across subtypes. Contingency test was done using GraphPad Prism v.8.4.3.

|  |  | All cohort | ER+ | HER2 | TNBC | p-values |
| --- | --- | --- | --- | --- | --- | --- |
| No. of patients |  | 75 | 31 | 20 | 24 |  |
| Age (n=75) | (Mean ± S.D) | 55.094 ± 11.77 | 56.16±12.99 | 50.52±9.46 | 54.75±11.96 | 0.7498 |
|  | Early (<50) | 31.08% (23) | 32.26% (10) | 35% (7) | 25% (6) |  |
| | Late (≥ 50) | 68% (51) | 67.74% (21) | 65% (13) | 75% (18) | |
| Menopausal status (n=65) | Pre | 23.44% (15) | 18.52% (5) | 29.41% (5) | 23.81% (5) | 0.7023 |
|  | Post | 76.56% (49) | 81.48% (22) | 70.59% (12) | 76.19% (16) |  |
| Grade (n=72) | Low (I/II) | 49.33% (37) | 77.42 % (24) | 44.44% (8) | 26.09% (6) | 0.0007 |
|  | High (III) | 45.33% (34) | 22.58% (7) | 55.56% (10) | 73.91% (17) |  |
| Tumor size (cT) (n=68) | T1 | 24% (18) | 40% (12) | 7.14% (1) | 25% (6) | 0.2752 |
|  | T2 | 58.67% (44) | 56.67% (17) | 85.71% (12) | 62.5% (15) |  |
|  | T3 | 5.33% (4) | 3.33% (1) | 7.14% (1) | 8.33% (2) |  |
|  | T4 | 1.33% (1) | 0 | 0 | 4.17% (1) |  |
| LVI (n=73) | Negative | 82.19% (60) | 80.65% (25) | 89.47% (17) | 78.26% (18) | 0.2658 |
|  | Positive | 17.81% (13) | 19.35% (6) | 10.53% (2) | 21.74% (5) |  |
| pT (primary tissue, no NACT) (n=39) | T0 | 5.13% (2) | 14.29% (2) | 0 | 0 | 0.4035 |
|  | T1 | 23.08% (9) | 28.57% (4) | 23.08% (3) | 16.67% (2) |  |
|  | T2 | 64.10% (25) | 42.86% (6) | 76.92% (10) | 75% (9) |  |
|  | T3 | 2.56% (1) | 7.14% (1) | 0 | 0 |  |
|  | T4 | 5.13% (2) | 7.14% (1) | 0 | 8.33% (1) |  |
| pN (primary tissue, no NACT) (n=39) | Negative | 69.23% (27) | 57.14% (8) | 76.92% (10) | 75% (9) | 0.4703 |
|  | Positive | 30.77% (12) | 42.86% (6) | 23.08% (3) | 25% (3) |  |
| Pathological Stage (primary tissue, no NACT) (n= 39) | Early(<IIB) | 66.67% (26) | 57.14% (8) | 76.92% (10) | 66.67% (8) | 0.5524 |
| | Late(≥IIB) | 33.33% (13) | 42.86% (6) | 23.08% (3) | 33.33% (4) | |
| NACT (n=74) | No | 60.81% (45) | 60% (18) | 65% (13) | 58.33% (14) | 0.897 |
|  | Yes | 39.19% (29) | 40% (12) | 35% (7) | 41.67% (10) |  |
| PCR status after NACT (n= 26) | pCR | 15.38% (4) | 9.09% (1) | 40% (2) | 10% (1) | 0.2364 |
|  | RD | 84.62% (22) | 90.91% (10) | 60% (3) | 90% (9) |  |
| Survival outcomes | No. followed-up | 68 | 27 | 19 | 23 |  |
|  | Median months | 29.97 | 35.5 | 34.77 | 29.6 |  |
|  | Follow-up in Months (Range) | 0.10-94.03 | 0.53-94.03 | 24.63-82.37 | 0.10-82.60 |  |
|  | # Recurred (local, distant) | 11 | 2 | 1 | 8 |  |
|  | # Death due to disease | 4 | 1 | 0 | 3 |  |

### NELFA expression and its association with survival outcomes

Immunohistochemistry for NELFA and NELFB was optimised using a standard protocol (Figure S5A-D)). Since the NELFA antibody showed a more robust and sharper IHC staining pattern compared to that of NELFB (Figure S5B-C), the cohort of 75 patient samples was stained and scored for NELFA expression, including percentage and intensity. Composite expression scores were computed by multiplying binned percent scores (0: 0% score, 1: 1-10%, 2: 11-50% and 3: 51-100%) with the intensity scores. Patients were classified into high (n = 37) or low (n = 38) NELFA expression categories based on the composite score cut-off from the ROC curve (Figure S6). Representative images of high and low NELFA are shown in Figure 3A.

**Figure 3.**
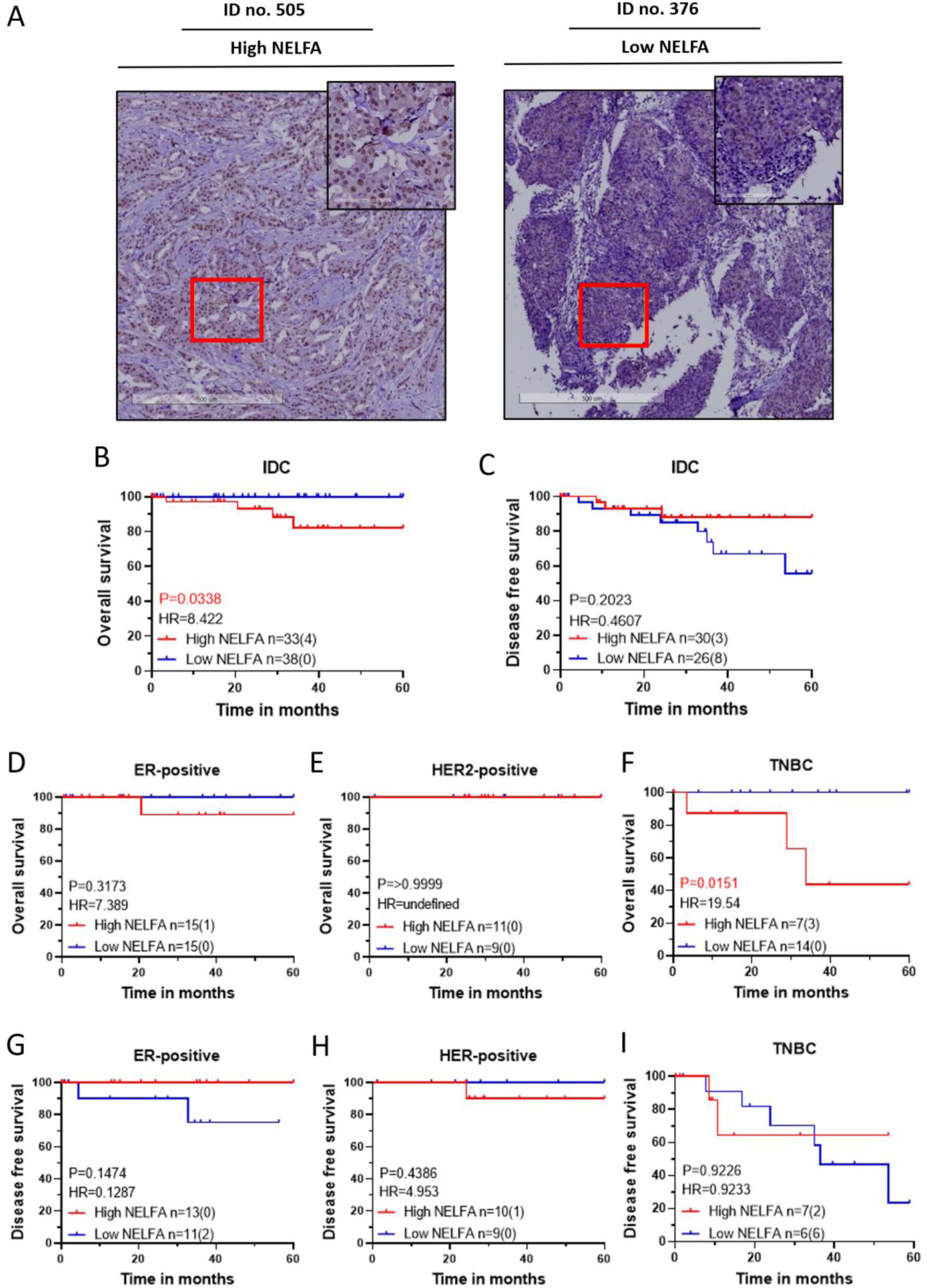
NELFA expression and its association with survival outcomes. NELFA expression and its association with patient survival outcomes. **(A)** Representative immunohistochemistry (IHC) images of invasive ductal carcinoma (IDC) tumours displaying high (#505) and low (#376) NELFA expression, with scale bars of 500 µm (overview) and 100 µm (zoomed region) included for reference. **(B-I)** Kaplan-Meier (KM) plots are based on the NELFA expression cohort. Statistical significances are computed using the log-rank test (Mantel-Cox) in GraphPad Prism 8.0.1 (244). For each KM Plot, the number of patients in each NELFA expression category is listed, along with the number of events in parentheses. **(B)** Overall Survival of the IDC cohort, **(C)** Disease-free survival of the IDC cohort, **(D-F)** Overall Survival of the molecular subtypes, and **(G-I)** Disease-free survival of the molecular subtypes.

The analysis of binned NELFA expression revealed no statistically significant association with the clinical features of the tumor at presentation (Table 6). Furthermore, the NELFA scores were evaluated for their association with survival outcomes. High NELFA expression was found to be significantly associated with poorer overall survival (Figure 3B), while low NELFA expression was associated with higher rates of recurrence, though not significant (Figure 3C).

**Table 6:** Association of NELFA expression with the clinical features of the IDC patients. IDC patient cohort was classified into High-NELFA and Low-NELFA groups based on the NELFA expression ROC curve with reference to the disease-free survival. A comparative assessment of High-NELFA versus Low-NELFA groups was performed using contingency analyses appropriate for each variable’s number of categorical levels, across clinical and pathological features. All statistical analyses were conducted using GraphPad Prism v.8.4.3.

**Table No.6: Association of NELFA expression with the clinical features of the IDC patients**
|  |  | All cohort | High NELFA | Low NELFA | p-values |
| --- | --- | --- | --- | --- | --- |
| No. of patients |  | 75 | 37 | 38 |  |
| Age (n=74) | (Mean ± S.D) | 55.094 ± 11.77 | 54.42 ± 11.66 | 55.74 ± 12.00 | 0.5501 |
|  | Early (<50) | 31.08% (23) | 27.03% (10) | 34.21% (13) |  |
| | Late (≥ 50) | 68% (51) | 70.27% (26) | 65.79% (25) | |
| Menopausal status (n=64) | Pre | 23.44% (15) | 21.62% (8) | 18.42% (7) | 0.6646 |
|  | Post | 76.56% (49) | 62.16% (23) | 68.42% (26) |  |
| Grade (n=71) | Low (I/II) | 49.33% (37) | 51.35% (19) | 47.37% (18) | 0.904 |
|  | High (III) | 45.33% (34) | 45.95% (17) | 44.74% (17) |  |
| Tumor size (cT) (n=67) | T1 | 24% (18) | 21.62% (8) | 26.32% (10) | 0.7575 |
|  | T2 | 58.67% (44) | 56.76% (21) | 60.53% (23) |  |
|  | T3 | 5.33% (4) | 5.4% (2) | 5.26% (2) |  |
|  | T4 | 1.33% (1) | 2.7% (1) | 0 |  |
| Node (cN) (n=71) | Negative | 29.33% (22) | 24.32% (9) | 34.21% (13) | 0.3436 |
|  | Positive | 65.33% (49) | 70.27% (26) | 60.53% (23) |  |
| pT (primary tissue, no NACT) (n=65) | T0 | 8% (6) | 10.81% (4) | 5.26% (2) | 0.7988 |
|  | T1 | 26.67% (20) | 29.73% (11) | 23.68% (9) |  |
|  | T2 | 45.33% (34) | 40.54% (15) | 50% (19) |  |
|  | T3 | 2.67% (2) | 2.7% (1) | 2.63% (1) |  |
|  | T4 | 4% (3) | 5.4% (2) | 2.63% (1) |  |
| pN (primary tissue, no NACT) (n=38) | Negative | 34.67% (26) | 32.43% (12) | 36.84% (14) | 0.4852 |
|  | Positive | 16% (12) | 18.92% (7) | 13.16% (5) |  |
| Pathological Stage (primary tissue, no NACT) (n= 38) | Early(<IIB) | 33.33% (25) | 29.73% (11) | 36.84% (14) | 0.305 |
| | Late(≥IIB) | 17.33% (13) | 21.62% (8) | 13.16% (5) | |
| NACT (n=73) | No | 60.27% (44) | 57.14% (20) | 63.16% (24) | 0.5998 |
|  | Yes | 39.73% (29) | 42.86% (15) | 36.84% (14) |  |
| PCR status after NACT (n= 26) | pCR | 13.79 (4) | 15.38% (2) | 15.38% (2) | >0.9999 |
|  | RD | 75.86% (22) | 84.62% (11) | 84.62% (11) |  |
| Subtype | ER+ | 41.33% (31) | 43.24% (16) | 39.47% (15) |  |
|  | HER2+ | 26.67% (20) | 29.73% (11) | 23.68% (9) |  |
|  | TNBC | 32.00% (24) | 27.03% (10) | 36.84% (14) |  |
| Survival outcomes | No. followed-up | 68 | 35.00 | 33.00 |  |
|  | Median months | 29.97 | 29.87 | 29.87 |  |
|  | Follow-up in Months (Range) | 0.10-94.03 | 0.13-94.03 | 0.10-82.60 |  |
|  | # Recurred (local, distant) | 11 | 3.00 | 8 |  |
|  | # Death due to disease | 4 | 4 | 0 |  |

Molecular Subtype-wise analysis further demonstrated that specifically the TNBC subtype showed a significant association with high NELFA expression and the worst overall survival (Figure 3F), but not with the ER-positive (Figure 3D) or HER2-positive subtypes (Figure 3E). For disease-free survival, the ER-positive subtype showed an inverse association with NELFA expression, approaching significance (Figure 3G), but not in HER2-positive (Figure 3H) or TNBC (Figure 3I) subtypes.

### YAP expression and its association with survival outcomes

The same breast cancer cohort tumor samples were stained for YAP by IHC and scored by a certified pathologist, using the same grading system as for NELFA expression. ROC curve was generated to determine the expression cut-off. Patients were classified as having high (n = 37) or low (n = 38) YAP expression. Representative images of high and low YAP are shown in Figure 4A. Clinical association with high and low YAP expression is shown in Table 7. Survival analysis revealed that high YAP expression was consistently associated with a poor prognosis across both overall and disease-free survival (Figure 4B-C).

**Figure 4.**
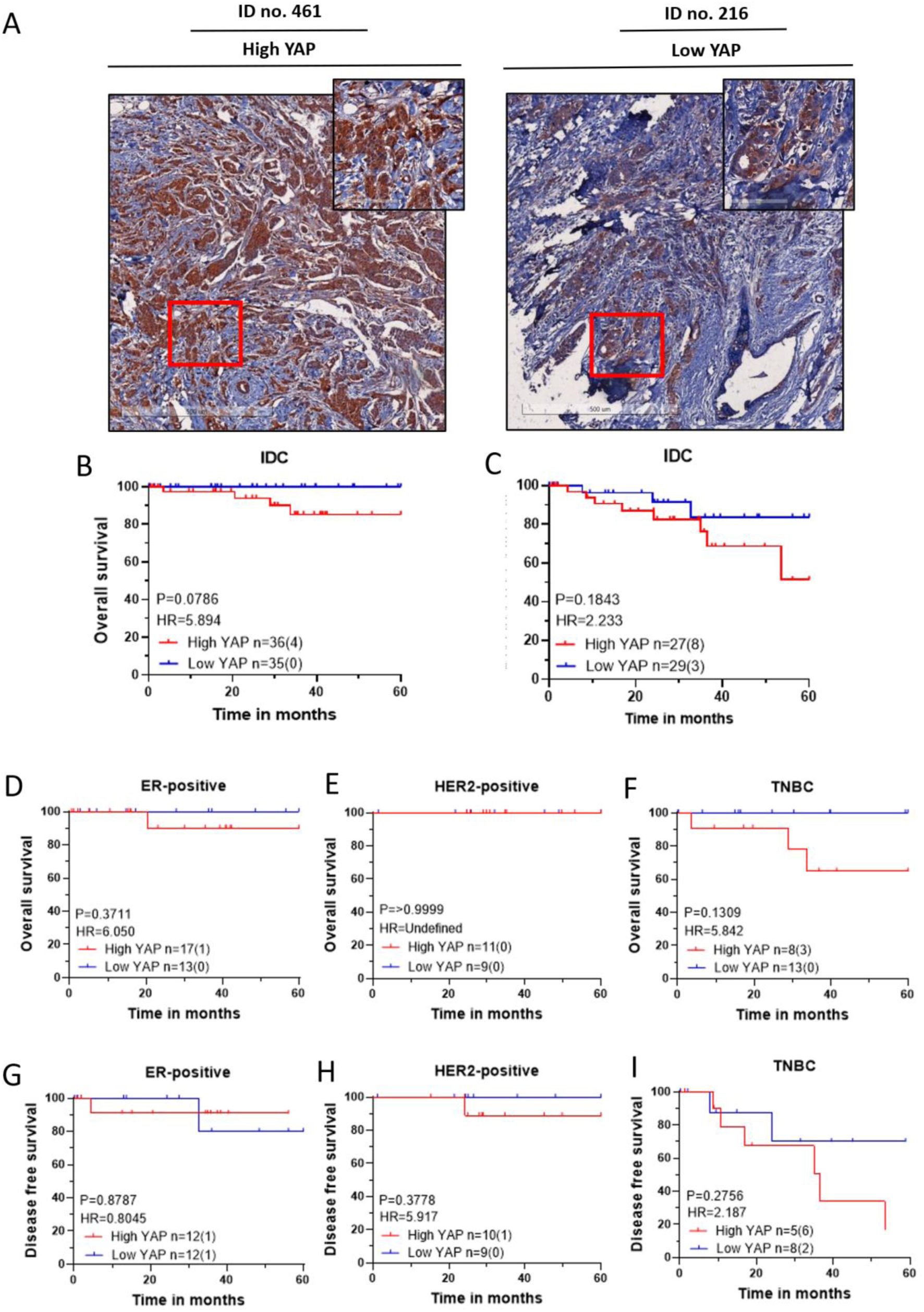
High YAP expression correlates with poor survival: **(A)** Representative immunohistochemistry (IHC) images of invasive ductal carcinoma (IDC) tumours displaying high (#499) and low (#216) YAP expression, with scale bars of 500 µm (overview) and 100 µm (zoomed region) included for reference. **(B-I)** Kaplan-Meier (KM) plots are based on the YAP expression cohort. Statistical significances are computed using the log-rank test (Mantel-Cox) in GraphPad Prism 8.0.1 (244). For each KM Plot, the number of patients in each YAP expression category is listed, along with the number of events in parentheses. **(B)** Overall Survival of the IDC cohort, **(C)** Disease-free survival of the IDC cohort, **(D-F)** Overall Survival of the molecular subtypes, and **(G-I)** Disease-free survival of the molecular subtypes.

**Table 7:** Association of YAP expression with the clinical features of the IDC patients. A cohort of IDC patients was categorized according to High-YAP and Low-YAP groups. It was based on the YAP expression ROC curve with reference to the disease-free survival. The distribution of clinical and pathological features and parameters between High-YAP and Low-YAP groups were evaluated using contingency analyses. Contingency test was done using GraphPad Prism v.8.4.3.

**Table No.7: Association of YAP expression with the clinical features of the IDC patients**
|  |  | All cohort | High YAP | Low YAP | p-values |
| --- | --- | --- | --- | --- | --- |
| No. of patients |  | 75 | 40 | 35 |  |
| Age (n=74) | (Mean ± S.D) | 55.094 ± 11.77 | 54.985 ± 11.99 | 55.094 ± 11.77 | 0.7748 |
|  | Early (<50) | 31.08% (23) | 32.50% (13) | 29.41% (10) |  |
| | Late (≥ 50) | 68% (51) | 67.50% (27) | 70.59% (24) | |
| Menopausal status (n=64) | Pre | 23.44% (15) | 20.00% (7) | 27.59% (8) | 0.4757 |
|  | Post | 76.56% (49) | 80.00% (28) | 72.41% (21) |  |
| Grade (n=71) | Low (I/II) | 49.33% (37) | 63.16% (24) | 39.39% (13) | 0.0456 |
|  | High (III) | 45.33% (34) | 36.84% (14) | 60.61% (20) |  |
| Tumor size (cT) (n=67) | T1 | 24% (18) | 29.41% (10) | 24.24% (8) | 0.5304 |
|  | T2 | 58.67% (44) | 64.71% (22) | 66.67% (22) |  |
|  | T3 | 5.33% (4) | 2.94% (1) | 9.09% (3) |  |
|  | T4 | 1.33% (1) | 2.94 (1) | 0 |  |
| Node (cN) (n=71) | Negative | 29.33% (22) | 21.62% (8) | 41.18% (14) | 0.0751 |
|  | Positive | 65.33% (49) | 78.38% (29) | 58.82% (20) |  |
| pT (primary tissue, no NACT) (n=65) | T0 | 8% (6) | 5.88% (2) | 12.90% (4) | 0.4157 |
|  | T1 | 26.67% (20) | 29.41% (10) | 32.26% (10) |  |
|  | T2 | 45.33% (34) | 58.82% (20) | 45.16% (14) |  |
|  | T3 | 2.67% (2) | 0 | 6.45% (2) |  |
|  | T4 | 4% (3) | 5.88% (2) | 3.23% 91) |  |
| pN (primary tissue, no NACT) (n=38) | Negative | 34.67% (26) | 70.00% (14) | 66.67% (12) | 0.8253 |
|  | Positive | 16% (12) | 30.00% (6) | 33.33% (6) |  |
| Pathological Stage (primary tissue, no NACT) (n= 38) | Early(<IIB) | 33.33% (25) | 65.00% (13) | 66.67% (12) | 0.9139 |
| | Late(≥IIB) | 17.33% (13) | 35.00% (7) | 33.33% (6) | |
| NACT (n=73) | No | 60.27% (44) | 61.54% (24) | 58.82% (20) | 0.8131 |
|  | Yes | 39.73% (29) | 38.46% (15) | 41.18% (14) |  |
| PCR status after NACT (n= 68) | pCR | 13.79 (4) | 15.38% (2) | 15.38% (2) | >0.9999 |
|  | RD | 75.86% (22) | 84.62% (11) | 84.62% (11) |  |
| Subtype | ER+ | 41.33% (31) | 45% (18) | 38.24% (13) | 0.7531 |
|  | HER2+ | 26.67% (20) | 27.5% (11) | 26.47% (9) |  |
|  | TNBC | 32.00% (24) | 27.5% (11) | 35.29% (12) |  |
| Survival outcomes | No. followed-up | 68 | 36 | 32 |  |
|  | Median months | 29.97 | 30.07 | 29.87 |  |
|  | Follow-up in Months (Range) | 0.10-94.03 | 0.53-82-.60 | 0.10-94.03 |  |
|  | # Recurred (local, distant) | 11 | 8 | 3 |  |
|  | # Death due to disease | 4 | 4 | 0 |  |

The subtype-wise analysis further confirmed that similar trend, irrespective of the molecular subtype (Figure 4D-I). Among these, TNBC showed the most separation, aligning with prior reports that YAP is frequently hyperactivated in TNBC and contributes directly to its aggressive phenotype [36]. These findings are consistent with the extensive literature establishing YAP as a driver of proliferation, survival, and metastasis in breast cancer [10].

### Association of NELFA expression in the context of YAP

To further assess the role of the interplay between the promoter-proximal pausing (PPP) component NELFA and YAP in breast cancer progression, we performed a survival analysis of NELFA expression levels in the context of YAP expression. Patients were stratified into four groups based on high or low expression of both YAP and NELFA. Representative images depicting these four groups are shown in Figure 5A. Clinical association with high and low NELFA plus YAP grouped expression is shown in Table no. 8.

**Figure 5.**
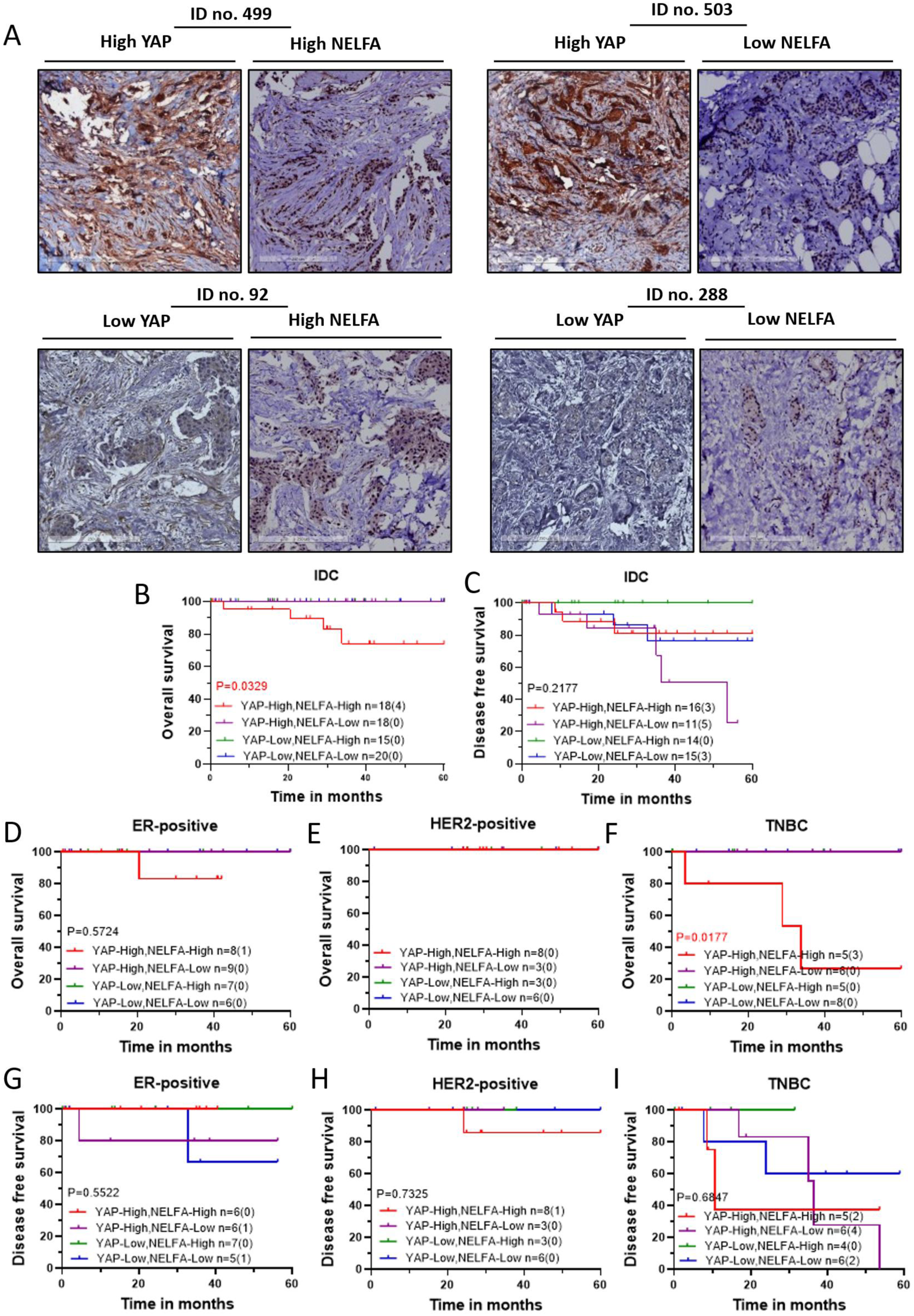
YAP and NELFA expression and its association with patient survival outcomes. **(A)** Representative immunohistochemistry (IHC) images of invasive ductal carcinoma (IDC) tumours displaying combination of high and low YAP expression with high and low NELFA. YAP-High, NELFA-High (#499); YAP-High, NELFA-Low (#503); YAP-Low, NELFA-High (#92); YAP-Low, NELFA-Low (#288), with scale bars of 250 µm (overview) included for reference. **(B-I)** Kaplan-Meier (KM) plots are based on the combination of YAP and NELFA expression cohort. Statistical significances are computed using the log-rank test (Mantel-Cox) in GraphPad Prism 8.0.1 (244). For each KM Plot, the number of patients in each YAP and NELFA expression category is listed, along with the number of events in parentheses. **(B)** Overall Survival of the IDC cohort, **(C)** Disease-free survival of the IDC cohort, **(D-F)** Overall Survival of the molecular subtypes, and **(G-I)** Disease-free survival of the molecular subtypes.

**Table 8:** Association of YAP and NELFA (combined) expression with the clinical features of the IDC patients. IDC samples were categorised into four combined-expression groups: High-YAP/High-NELFA, High-YAP/Low-NELFA, Low-YAP/High-NELFA, and Low-YAP/Low-NELFA. The table outlines demographic, clinical, and pathological variables, including age at diagnosis, menopausal status, tumour grade, radiological and pathological tumour dimensions, disease stage and treatment details. Statistical analyses were performed using GraphPad Prism v.8.4.3.

**Table No.8: Association of YAP and NELFA (combined) expression with the clinical features of the IDC patients.**
|  |  | All cohort | High YAP and High NELFA | High YAP and Low NELFA | Low YAP and High NELFA | Low YAP and Low NELFA | p-values |
| --- | --- | --- | --- | --- | --- | --- | --- |
| No. of patients |  | 75 | 22 | 18 | 15 | 20 |  |
| Age (n=74) | (Mean ± S.D) | 55.094 ± 11.77 | 54.985 ± 11.99 | 54.473 ± 12.74 | 55.094 ± 11.77 | 55.043 ± 11.98 | 0.8493 |
|  | Early (<50) | 31.08% (23) | 31.82% (7) | 33.33% (6) | 21.43% (3) | 35% (7) |  |
| | Late (≥ 50) | 68% (51) | 68.18% (15) | 66.67% (12) | 78.57% (11) | 65% (13) | |
| Menopausal status (n=64) | Pre | 23.44% (15) | 26.32% (5) | 12.5% (2) | 25% (3) | 29.41% (5) | 0.6802 |
|  | Post | 76.56% (49) | 73.68% (14) | 87.5% (14) | 75% (9) | 70.59% (12) |  |
| Grade (n=71) | Low (I/II) | 49.33% (37) | 57.143% (12) | 70.59% (12) | 46.67% (7) | 33.33% (6) | 0.1537 |
|  | High (III) | 45.33% (34) | 42.86% (9) | 29.41% (5) | 53.33% (8) | 66.67% (12) |  |
| Tumor size (cT) (n=67) | T1 | 24% (18) | 23.53% (4) | 35.29% (6) | 26.67% (4) | 22.22% (4) | 0.7031 |
|  | T2 | 58.67% (44) | 70.59% (12) | 58.82% (12) | 60% (9) | 72.22% (13) |  |
|  | T3 | 5.33% (4) | 0 | 5.88% (1) | 13.33% (2) | 5.56% (1) |  |
|  | T4 | 1.33% (1) | 5.88% (1) | 0 | 0 | 0 |  |
| Node (cN) (n=71) | Negative | 29.33% (22) | 20% (4) | 23.53% (4) | 33.33% (5) | 47.37% (9) | 0.2621 |
|  | Positive | 65.33% (49) | 80% (16) | 76.47% (13) | 66.67% (10) | 52.63% (10) |  |
| pT (primary tissue, no NACT) (n=65) | T0 | 8% (6) | 5.26% (1) | 6.67% (1) | 21.23% (3) | 5.88% (1) | 0.7435 |
|  | T1 | 26.67% (20) | 36.84% (7) | 20% (3) | 28.57% (4) | 35.29% (6) |  |
|  | T2 | 45.33% (34) | 52.63% (10) | 66.67% (10) | 35.71% (5) | 52.94% (9) |  |
|  | T3 | 2.67% (2) | 0 | 0 | 7.14% (1) | 5.88% (1) |  |
|  | T4 | 4% (3) | 5.26% (1) | 6.67% (1) | 7.14% (1) | 0 |  |
| pN (primary tissue, no NACT) (n=38) | Negative | 34.67% (26) | 75% (9) | 62.5% (5) | 42.86% (3) | 81.82% (9) | 0.3338 |
|  | Positive | 16% (12) | 25% (3) | 37.5% (3) | 57.14% (4) | 18.18% (2) |  |
| Pathological Stage (primary tissue, no NACT) (n= 38) | Early(<IIB) | 33.33% (25) | 66.67% (8) | 62.50% (5) | 42.86% (3) | 81.82% (9) | 0.4019 |
| | Late(≥IIB) | 17.33% (13) | 33.33% (4) | 37.50% (3) | 57.14% (4) | 18.18% (2) | |
| NACT (n=73) | No | 60.27% (44) | 61.90% (13) | 61.11% (11) | 50% (7) | 65% (13) | 0.8417 |
|  | Yes | 39.73% (29) | 38.09% (8) | 38.88% (7) | 50% (7) | 35% (7) |  |
| PCR status after NACT (n= 68) | pCR | 13.79 (4) | 16.66% (1) | 14.28% (1) | 14.28% (1) | 16.66% (1) | 0.9988 |
|  | RD | 75.86% (22) | 83.33% (5) | 85.71% (6) | 85.71% (6) | 83.33% (5) |  |
| Subtype (n=75) | ER+ | 41.33% (31) | 40.90% (9) | 50% (9) | 46.66% (7) | 30% (6) | 0.6975 |
|  | HER2+ | 26.67% (20) | 36.36% (8) | 16.66% (3) | 20% (3) | 30% (6) |  |
|  | TNBC | 32.00% (24) | 22.72% (5) | 33.33% (6) | 33.33% (5) | 40% (8) |  |
| Survival outcomes | No. followed-up | 68 | 22 | 14 | 13 | 19 |  |
|  | Median months | 29.97 | 30.7 | 34.77 | 29.87 | 29.87 |  |
|  | Follow-up in Months (Range) | 0.10-94.03 | 0.53-69.40 | 1.10-82.60 | 0.13-94.03 | 0.10-76.97 |  |
|  | # Recurred (local, distant) | 11 | 3 | 5 | 0 | 3 |  |
|  | # Death due to disease | 4 | 4 | 0 | 0 | 0 |  |

Patients with high YAP and high NELFA expression in the primary tumors showed significantly worse outcomes for overall survival (Figure 5B). While high YAP and low NELFA expression are associated with worse disease-free survival outcomes, though not significantly so (Figure 5C).

The association of high YAP and high NELFA expression with the overall survival was once again significant in the TNBC subtype (Figure 5F) but not for the ER or HER2 positive subtype (Figure 5E-H). For disease-free survival, a strong separation was observed in the outcomes for the four categories of TNBC subtype again (Figure 5I), but not for the other two subtypes (Figure 5G-H).

### Breast Cancer cohort from TCGA and its association with NELFA and YAP expression

To validate the correlation observed in the small cohort of breast cancer patient samples, the breast cancer cohort from the TCGA database was analyzed. A cohort of IDC (n=415) with associated clinical metadata and RNA-seq expression data was downloaded from cBioPortal. No significant association was observed for mRNA expression of YAP, NELFA, or YAP + NELFA expression with overall or disease-free survival (Figure S7). Subtype-wise analysis for the association revealed no significant association with survival outcomes for NELFA RNA expression (Figure 6A) or for YAP RNA expression (Figure 6B), except that the HER2-positive subtype showed a significant association with the worst outcome when YAP mRNA had high expression. Analysis of NELFA expression within a high YAP expression background revealed a significant association between low NELFA expression and worse overall survival, specifically for the HER2-positive subtype (Figure 6C). For disease-free survival, a similar trend was observed across all subtypes; however, none were statistically significant (Figure 6C).

**Figure 6.**
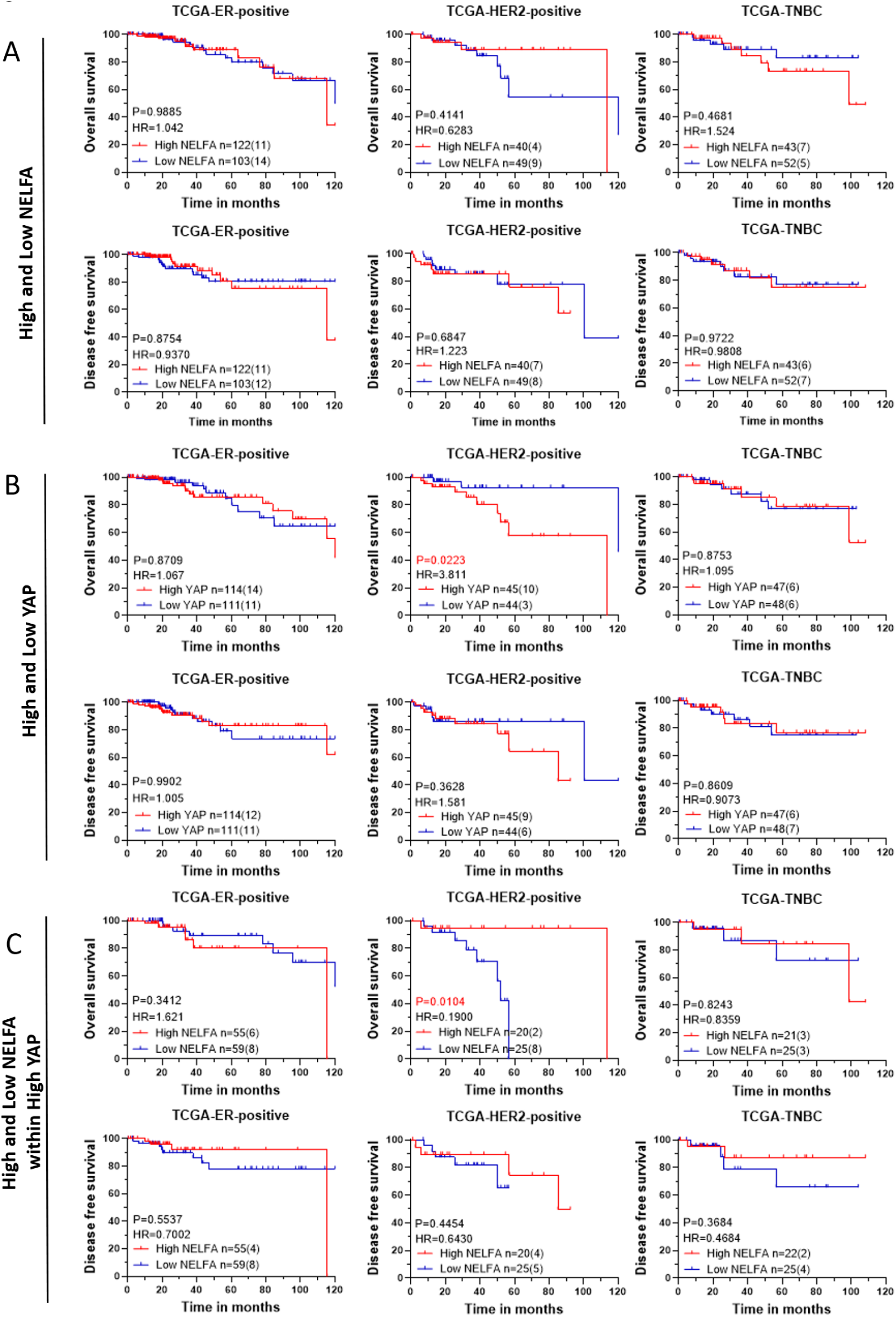
Overall survival and disease-free survival in IDC breast cancer cohort of TCGA. **(A-C)** KM plots of Breast cancer patients divided based expression of individual genes NELFA, YAP, and NELFA within High YAP expression of BC (Breast Cancer) IDC cohort. Statistical significances are computed using the log-rank test (Mantel-Cox) in GraphPad Prism 8.0.1 (244). **(A)** Overall and disease-free survival of NELFA expression for molecular subtypes of BC **(B)** Overall and disease-free survival of YAP expression for molecular subtypes of BC **(C)** Overall and disease-free survival of NELFA expression within high YAP for molecular subtypes of BC

## Discussion

Gene expression is a highly regulated process, with promoter-proximal pausing (PPP) representing one of the checkpoints [37–41]. Our *in vivo* screen in *Drosophila* [6] revealed that 7SK snRNP and NELF-A, components of the PPP complex, were part of the significant genes that enhanced *Yki*-driven hyperproliferation [5]. Building on this, we investigated whether the PPP complex-mediated regulation of YAP transcription is conserved in the mammalian system. Our findings revealed that silencing 7SK snRNP components MePCE and HEXIM1/2 in the mammalian cell line HEK293T did not alter YAP target gene expression, whereas depletion of NELFA did. NELFA-mediated regulation of YAP–target expression was observed in an independent mammalian cell line: MDA-MB-231, a breast cancer cell line. These findings indicate that NELF-A, but not the broader 7SK snRNP complex, selectively restrains YAP-mediated transcription in the mammalian system.

An exploratory whole-transcriptome analysis after NELFA knockdown in MDA-MB-231 revealed widespread transcriptional reprogramming, with the most significant alteration being a YAP target gene signature. Enrichment of Tumor Growth Factor (TGF) signaling, which is a known upstream activator of YAP and EMT-related pathways [42,43], supports a functional link between NELF-A loss and activation of YAP-mediated transcription. Furthermore, TF enrichment analysis of NELFA-regulated genes identified TEAD4, the canonical YAP partner, and CJUN, a component of the AP-1 complexes [32,44], among the top hits, indicating the involvement of the YAP-TEAD-AP-1 oncogenic axis. Previous studies have established that the NELF complex plays a critical role in transcriptional regulation in breast cancer cells, participating in canonical biological programs such as the Cell cycle, Proliferation, and EMT, among others [33,34] When the genes perturbed by NELF components in the breast cancer cell lines from these studies were compared with those from this study in MDA-MB-231, a very minimal overlap was observed, indicating cell line-specific regulation of NELF components.

To dissect functional crosstalk between NELFA and YAP-mediated transcription, we developed a log2 fold-change-based classification of co-regulated genes across siNELF-A, siYAP, and double-knockdown conditions. Four distinct functions were identified with additive, antagonistic, and cooperative regulation: 1. genes activated by YAP but repressed by NELFA, 2. Genes activated by NELF-A but suppressed by YAP, 3. Co-repressed, and 4. co-activated genes. Independent STRING analysis of these four categories revealed distinct biologically relevant modules.

Clinical analyses in an Indian breast cancer cohort further confirmed the role of the NELFA-YAP axis in patient survival outcomes. It also underscored the context-dependent role of NELFA. High NELFA correlated significantly with poorer overall survival and showed an opposite but not significant association with disease-free survival (DFS). Notably, in the context of high YAP expression, low NELFA exhibited the worst DFS outcomes, particularly in the TNBC subtype, mirroring our cell-line observations where NELF-A loss showed overexpression of YAP target genes involved in oncogenesis and EMT. Distinct outcomes associated with NELFA expression, when assessed independently or in the context of YAP expression, emphasize the context-specific role of NELFA.

The breast cancer patient cohort from the TCGA dataset was analyzed as an independent cohort to investigate the association between NELFA expression and patient outcomes. NELFA expression at the mRNA level did not show any specific association with patient outcomes, but when assessed in the context of High-YAP expression, patients with low NELFA expression showed a trend towards worse recurrence outcome and significantly worse survival outcomes. Thus, analysis of the TCGA cohort reinforced the tumor-suppressive dimension of NELFA specifically in the context of high YAP expression.

Collectively, our data indicate that NELFA regulates YAP-mediated transcription in the mammalian system, and the PPP-YAP axis is also conserved in this system. Furthermore, we identified a global network of genes regulated by NELFA and, consequently, the PPP complex. Our clinical data analysis for NELFA-YAP interaction in the breast cancer context revealed dual functions for NELFA, one as an oncogene when assessed independently and one as a tumor suppressor in the context of high-YAP expression.

## Author Contribution

BV, AA performed all the experiments. RK analyzed RNA sequencing data. KS supported cell culture experiments. CK, LSS, and MK supervised the study, provided resources, and funding. BV, AA, RK, and MK drafted the manuscript. MK conceptualized and supervised the experimentation and the data analysis. All authors read and approved the final version of the manuscript.

## Acknowledgements

The core facility at Indian Institute of Science Education and Research is acknowledged for providing quantification and imaging systems. Prashanti Cancer Care Mission biobank for providing the patient samples. Quantitative Pathology Imaging System Mantra^TM^ and Multimode Plate Reader EnSight^TM^ at Revvity IISER Pune Centre for Excellence, formerly, Perkin Elmer-IISER Pune Centre for Excellence, hosted at IISER Pune. BV would like to acknowledge Vaishnavi Jadhav for rectifying the CRISPR experiments and Anuvind Pramod for helping to download TCGA data from cBioPortal. BV would also like to thank all members of the Tumor Microenvironment lab for their continuous input and support.

## Bibliography

1 Zanconato F, Cordenonsi M, Piccolo S. YAP/TAZ at the Roots of Cancer. Cancer Cell. 2016;29(6):783–803.

2 Zhao B, Ye X, Yu J, Li L, Li W, Li S, et al. TEAD mediates YAP-dependent gene induction and growth control. Genes Dev. 2008;22(14):1962–1971.

3 Boopathy GTK, Hong W. Role of Hippo Pathway-YAP/TAZ Signaling in Angiogenesis. Front Cell Dev Biol. 2019;7:49.

4 Szulzewsky F, Holland EC, Vasioukhin V. YAP1 and its fusion proteins in cancer initiation, progression and therapeutic resistance. Dev Biol. 2021;475:205–221.

5 Nagarkar S, Wasnik R, Govada P, Cohen S, Shashidhara LS. Promoter Proximal Pausing Limits Tumorous Growth Induced by the Yki Transcription Factor in *Drosophila*. Genetics. 2020;216(1):67–77.

6 Groth C, Vaid P, Khatpe A, Prashali N, Ahiya A, Andrejeva D, et al. Genome-Wide Screen for Context-Dependent Tumor Suppressors Identified Using in Vivo Models for Neoplasia in *Drosophila*. G3 Genes|Genomes|Genetics. 2020;10(9):2999–3008.

7 Core L, Adelman K. Promoter-proximal pausing of RNA polymerase II: a nexus of gene regulation. Genes Dev. 2019;33(15–16):960–982.

8 Yamaguchi Y, Inukai N, Narita T, Wada T, Handa H. Evidence that Negative Elongation Factor Represses Transcription Elongation through Binding to a DRB Sensitivity-Inducing Factor/RNA Polymerase II Complex and RNA. Mol Cell Biol. 2002;22(9):2918–2927.

9 Vos SM, Farnung L, Boehning M, Wigge C, Linden A, Urlaub H, et al. Structure of activated transcription complex Pol II–DSIF–PAF–SPT6. Nature. 2018;560(7720):607–612.

10 Thompson BJ. YAP/TAZ: Drivers of Tumor Growth, Metastasis, and Resistance to Therapy. BioEssays. 2020;42(5).

11 Dobin A, Davis CA, Schlesinger F, Drenkow J, Zaleski C, Jha S, et al. STAR: Ultrafast universal RNA-seq aligner. Bioinformatics. 2013;29(1):15–21.

12 Deluca DS, Levin JZ, Sivachenko A, Fennell T, Nazaire MD, Williams C, et al. RNA-SeQC: RNA-seq metrics for quality control and process optimization. Bioinformatics. 2012;28(11):1530–1532.

13 Wang L, Wang S, Li W. RSeQC: Quality control of RNA-seq experiments. Bioinformatics. 2012;28(16):2184–2185.

14 Ewels P, Magnusson M, Lundin S, Käller M. MultiQC: Summarize analysis results for multiple tools and samples in a single report. Bioinformatics. 2016;32(19):3047–3048.

15 Liao Y, Smyth GK, Shi W. FeatureCounts: An efficient general purpose program for assigning sequence reads to genomic features. Bioinformatics. 2014;30(7):923–930.

16 Love MI, Huber W, Anders S. Moderated estimation of fold change and dispersion for RNA-seq data with DESeq2. Genome Biol. 2014;15(12).

17 Yu G, Wang LG, Han Y, He QY. ClusterProfiler: An R package for comparing biological themes among gene clusters. OMICS. 2012;16(5):284–287.

18 Korotkevich G, Sukhov V, Budin N, Shpak B, Artyomov MN, Sergushichev A. Fast gene set enrichment analysis. 2016.

19 Dolgalev I. msigdbr: MSigDB Gene Sets for Multiple Organisms in a Tidy Data Format. 2025.

20 Carlson M. org.Hs.eg.db: Genome wide annotation for Human. 2025.

21 Liberzon A, Birger C, Thorvaldsdóttir H, Ghandi M, Mesirov JP, Tamayo P. The Molecular Signatures Database Hallmark Gene Set Collection. Cell Syst. 2015;1(6):417–425.

22 Gao CH, Yu G, Cai P. ggVennDiagram: An Intuitive, Easy-to-Use, and Highly Customizable R Package to Generate Venn Diagram. Front Genet. 2021;12.

23 Krassowski M. ComplexUpset: Create Complex UpSet Plots Using ggplot2 Components. 2021.

24 Chen EY, Tan CM, Kou Y, Duan Q, Wang Z, Meirelles GV, et al. Enrichr: interactive and collaborative HTML5 gene list enrichment analysis tool. BMC Bioinformatics. 2013;14:128.

25 Wickham H, Chang W, Henry L, Pedersen TL, Takahashi K, Wilke C. ggplot2: Create Elegant Data Visualisations Using the Grammar of Graphics. 2025.

26 Cheng B, Price DH. Properties of RNA Polymerase II Elongation Complexes Before and After the P-TEFb-mediated Transition into Productive Elongation. Journal of Biological Chemistry. 2007;282(30):21901–21912.

27 Wang J, Rojas P, Mao J, Mustè Sadurnì M, Garnier O, Xiao S, et al. Persistence of RNA transcription during DNA replication delays duplication of transcription start sites until G2/M. Cell Rep. 2021;34(7):108759.

28 Venkatasubramanian G, Kelkar DA, Mandal S, Jolly MK, Kulkarni M. Analysis of Yes-Associated Protein-1 (YAP1) Target Gene Signature to Predict Progressive Breast Cancer. J Clin Med. 2022;11(7):1947.

29 Peterlin BM, Brogie JE, Price DH. 7SK snRNA: a noncoding RNA that plays a major role in regulating eukaryotic transcription. Wiley Interdiscip Rev RNA. 2012;3(1):92–103.

30 Subramanian A, Tamayo P, Mootha VK, Mukherjee S, Ebert BL, Gillette MA, et al. Gene set enrichment analysis: A knowledge-based approach for interpreting genome-wide expression profiles. Proc Natl Acad Sci USA. 2005;102(43):15545–15550.

31 Cordenonsi M, Zanconato F, Azzolin L, Forcato M, Rosato A, Frasson C, et al. The hippo transducer TAZ confers cancer stem cell-related traits on breast cancer cells. Cell. 2011;147(4):759–772.

32 Zanconato F, Forcato M, Battilana G, Azzolin L, Quaranta E, Bodega B, et al. Genome-wide association between YAP/TAZ/TEAD and AP-1 at enhancers drives oncogenic growth. Nat Cell Biol. 2015;17(9):1218–1227.

33 Sun J, Li R. Human negative elongation factor activates transcription and regulates alternative transcription initiation. Journal of Biological Chemistry. 2010;285(9):6443–6452.

34 Zhang J, Hu Z, Chung HH, Tian Y, Lau KW, Ser Z, et al. Dependency of NELF-E-SLUG-KAT2B epigenetic axis in breast cancer carcinogenesis. Nat Commun. 2023;14(1).

35 Szklarczyk D, Gable AL, Nastou KC, Lyon D, Kirsch R, Pyysalo S, et al. The STRING database in 2021: Customizable protein-protein networks, and functional characterization of user-uploaded gene/measurement sets. Nucleic Acids Res. 2021;49(D1):D605–D612.

36 Yu B, Su J, Shi Q, Liu Q, Ma J, Ru G, et al. KMT5A-methylated SNIP1 promotes triple-negative breast cancer metastasis by activating YAP signaling. Nat Commun. 2022;13(1):2192.

37 Gilmour DS, Lis JT. RNA polymerase II interacts with the promoter region of the noninduced hsp70 gene in Drosophila melanogaster cells. Mol Cell Biol. 1986;6(11):3984–9.

38 Kwak H, Fuda NJ, Core LJ, Lis JT. Precise maps of RNA polymerase reveal how promoters direct initiation and pausing. Science. 2013;339(6122):950–3.

39 Jonkers I, Lis JT. Getting up to speed with transcription elongation by RNA polymerase II. Nat Rev Mol Cell Biol. 2015;16(3):167–77.

40 Noe Gonzalez M, Blears D, Svejstrup JQ. Causes and consequences of RNA polymerase II stalling during transcript elongation. Nat Rev Mol Cell Biol. 2021;22(1):3–21.

41 Rodríguez-Molina JB, West S, Passmore LA. Knowing when to stop: Transcription termination on protein-coding genes by eukaryotic RNAPII. Mol Cell. 2023;83(3):404–415.

42 Szeto SG, Narimatsu M, Lu M, He X, Sidiqi AM, Tolosa MF, et al. YAP/TAZ Are Mechanoregulators of TGF-β-Smad Signaling and Renal Fibrogenesis. Journal of the American Society of Nephrology. 2016;27(10):3117–3128.

43 Zhang T, He X, Caldwell L, Goru SK, Ulloa Severino L, Tolosa MF, et al. NUAK1 promotes organ fibrosis via YAP and TGF-β/SMAD signaling. Sci Transl Med. 2022;14(637).

44 Meng Q, Xia Y. c-Jun, at the crossroad of the signaling network. Protein Cell. 2011;2(11):889–898.

